# Thalamic state influences timing precision in the thalamocortical circuit

**DOI:** 10.1101/439778

**Authors:** Clarissa J Whitmire, Yi Juin Liew, Garrett B Stanley

## Abstract

Sensory signals from the outside world are transduced at the periphery, passing through thalamus before reaching cortex, ultimately giving rise to the sensory representations that enable us to perceive the world. The thalamocortical circuit is particularly sensitive to the temporal precision of thalamic spiking due to highly convergent synaptic connectivity. Thalamic neurons can exhibit burst and tonic modes of firing that strongly influence timing within the thalamus. The impact of these changes in thalamic state on sensory encoding in the cortex, however, remains unclear. Here, we investigated the role of thalamic state on timing in the thalamocortical circuit of the vibrissa pathway in the anesthetized rat. We optogenetically hyperpolarized thalamus while recording single unit activity in both thalamus and cortex. Tonic spike triggered analysis revealed temporally precise thalamic spiking that was locked to white-noise sensory stimuli, while thalamic burst spiking was associated with a loss in stimulus-locked temporal precision. These thalamic state dependent changes propagated to cortex such that the cortical timing precision was diminished during the hyperpolarized (burst biased) thalamic state. While still sensory driven, the cortical neurons became significantly less precisely locked to the white-noise stimulus. The results here suggest that tonic thalamic spiking is more temporally precise than burst firing, which leads to distinct differences in sensory information representation at the level of both the thalamus and the cortex, as assessed using spike triggered analysis. This difference in spike timing precision enables a dynamic encoding scheme for sensory information as a function of thalamic state.

**New and Noteworthy:** The majority of sensory signals are transmitted through the thalamus. There is growing evidence of complex thalamic gating through coordinated firing modes that have a strong impact on cortical sensory representations. Optogenetic hyperpolarization of thalamus pushed it into burst firing that disrupted precise time-locked sensory signaling, with a direct impact on the downstream cortical encoding, setting the stage for a timing-based thalamic gate of sensory signaling.

## Introduction

Sensory thalamus plays a critical role in gating information flow from sensors in the periphery to cortex, ultimately shaping how we perceive the world. Importantly, thalamic gating properties are not static, but instead vary dynamically through a range of modulatory mechanisms, including local membrane and synaptic properties (Wolfart et al., 2005), stimulus history (Whitmire et al., 2016), and neuromodulatory inputs (Castro-alamancos, 2002; Mease et al., 2014). Although arising from different mechanisms, these modulatory inputs have the net effect of altering the baseline membrane polarization level in the thalamus, referred to here as “thalamic state”, which plays an important role in determining the encoding properties of the thalamic neurons that serve as primary inputs to sensory cortex. Perhaps most prominently, modulation of the baseline membrane potential in thalamic neurons enables distinct tonic and burst firing modes due to the selective engagement of low threshold calcium channels during prolonged hyperpolarization (Suzuki and Rogawski, 1989). It has long been posited that these two firing modes could dynamically control information processing (Sherman 2001), but the precise way in which this could happen has remained speculative and the way in which cortical coding properties are shaped is unknown.

Although the large majority of studies of T-type calcium channel bursts in thalamus have been focused on the underlying detailed biophysical mechanisms enabled by brain slice recordings, there have been a number of investigations of the intact circuitry in-vivo. At the thalamocortical synapse, in-vivo studies have shown that spontaneous burst spikes are more effective at driving cortical spiking (Swadlow and Gusev, 2001) and evoke larger cortical depolarizations (Bruno and Sakmann, 2006) than tonic spikes. The in-vivo properties of burst and tonic spiking have been explored perhaps most extensively in the visual pathway (Alitto et al., 2005; Denning and Reinagel, 2005; Lesica and Stanley, 2004; Reinagel et al., 1999; Wang et al., 2007), with burst firing shown to be reliably elicited across trials in response to visual stimulation (Lesica and Stanley, 2004; Martinez-Conde et al., 2002; Wang et al., 2007), associated with an “all-or-none” type of response to facilitates detection of changes in the visual scene (Lesica and Stanley, 2004) consistent with mechanisms that would serve as a “wake-up call” to cortex (Sherman, 2001). Furthermore, although historically controversial, it has been shown in a number of studies that thalamic bursting is not just observed in sleep states or under anesthesia, but is present, albeit reduced, in the awake brain (Borden et al., 2019; Guido and Weyand, 1995; Whitmire et al., 2016). However, the implication for downstream cortical encoding remains elusive because of the complexity of the thalamocortical circuitry. It has been estimated that 50-100 thalamic neurons converge as the primary drivers of a single cortical neuron (Bruno and Sakmann, 2006), where the concerted effort of a relatively large number of synaptic inputs is needed to drive suprathreshold cortical activity. Without a mechanism to manipulate the population activity of thalamic neurons converging on a common cortical target independent of the sensory drive, the role of tonic/burst firing in driving downstream cortical activity remains elusive.

To address this, we used optogenetic manipulation of thalamic state to systematically bias the thalamic population towards a burst firing regime to quantify the role of thalamic state on precise timing of spiking activity in the thalamocortical circuit of the rodent whisker pathway using a two-stage, linear-nonlinear framework. The first stage represents the sensory feature selectivity, and the second stage represents the overall sensitivity of the input-output relationship (Estebanez et al., 2012; Petersen et al., 2008; Ramirez et al., 2014). This characterization was performed for neurons recorded extracellularly in the ventro posterior-medial (VPm) thalamus and in primary somatosensory cortex (S1) in the fentanyl-anesthetized rat, for both first order (spike-triggered average) and higher order (spike-triggered covariance) selectivity. For thalamic neurons, we found that tonic spiking was associated with whisker-stimulus feature selectivity consistent with previous findings (Petersen et al., 2008). However, analysis of burst firing suggested that while bursting activity was clearly sensory driven, the spike-triggered analysis revealed a lack of precise stimulus-locked spiking in response to white-noise whisker stimulation. This was further confirmed using optogenetic hyperpolarization of VPm to switch the thalamus into a burst mode. In the cortical neurons, when the thalamus was dominated by tonic firing, the cortical neurons exhibited similar feature selectivity as observed in VPm. However, when the thalamus was optogenetically hyperpolarized, the spike-triggered analysis revealed a reduction in precise stimulus-locked spiking in S1 units in response to the white-noise whisker stimulus, yet maintained a consistent overall stimulus-driven firing rate. Given the sensitivity of the cortex to precise timing of thalamic projection neurons, these results suggest that shifts in thalamic state disrupt precise timing of thalamic inputs to cortex that are compensated for by the potency of the thalamic bursts, setting the stage for a timing-based gating of information flow to cortex.

## Methods

### Experimental Procedures

#### Acute Surgery

All procedures were approved by the Georgia Institute of Technology Institutional Animal Care and Use Committee and were in agreement with guidelines established by the National Institutes of Health. 19 female albino rats (Sprague-Dawley, 250-300g) were anesthetized intravenously using a fentanyl cocktail (fentanyl (5 µg/kg), midazolam (2 mg/kg), dexmedetomidine (150 µg/kg)). A craniotomy was performed over VPm (2-4 mm caudal to bregma, 1.5-3.5 mm lateral to the midline), and in a subset of animals, a second craniotomy was performed over S1 (1-3 mm caudal to bregma, 4.5-6 mm lateral to the midline). At the termination of the experiment, the animal was euthanized with an overdose of sodium pentobarbital (euthasol, 0.5 mL at 390 mg/mL). All optogenetically transfected animals that underwent cortical probe recordings were perfused and their brains were imaged for verification of opsin location and cortical probe location.

#### Electrophysiology

Tungsten microelectrodes were lowered into the thalamus (depth: 4.5-6 mm) using a micropositioner (Kopf, Luigs-Neumann). Multielectrode probes (A1×32-10mm-50-177, NeuroNexus) were lowered perpendicular to S1 (45° relative to vertical; depth: 2 mm). The topographic location of the electrode was identified through manual stimulation of the whisker pad. Upon identification of the primary whisker for the recorded unit(s), the primary whisker was threaded into the galvo motor to permit stimulation of a single whisker.

#### Sensory Stimulus

Mechanical whisker stimulation was delivered using a precisely controlled galvo motor (Cambridge Technologies, custom Matlab software). The mechanical stimulus applied to the whisker in the rostral-caudal direction consisted of sensory white-noise (low pass filtered at 200 Hz, standard deviation of the noise was 0.6° or 223°/s). Feedback from the whisker stimulator were used for further spike triggered analysis across all units (down sampled to 4.88 kHz).

#### Optogenetics surgeries

All surgical procedures followed sterile protocol. A small craniotomy was made above VPm (3 mm lateral, 3 mm caudal to bregma). A 10 µL syringe (Neuros Syringe, Hamilton, Inc) filled with the virus (rAAV5-CamKIIa-Jaws-KGC-GFP-ER2 or rAAV5-CamKIIa-eNpHR3.0-EYFP, UNC Viral Vector Core Services) was lowered to depth of 5.2 mm before injecting 1 µL of virus at a rate of 0.2 µL/min (iSi system, Stoelting). The syringe remained in place for five minutes after the injection was complete to allow the virus to diffuse. Opsin expression was fully realized at 2-3 weeks post-surgery.

#### Optogenetic Stimulus

Optical manipulation was administered with a controlled pulse of light through a custom optrode consisting of an optical fiber (200µm diameter; Thorlabs) and an electrode (Tungsten microelectrode; FHC) that was lowered into the VPm. Upon identifying a whisker sensitive cell, light sensitivity was assessed by the post-inhibitory rebound spiking response using a train of 250 millisecond light pulses (λ = 590 or 617nm for Halorhodopsin and Jaws, respectively). The whisker was then stimulated without (baseline) and with (hyperpolarized) light provided directly to the thalamus (50 mW/mm^2^). Optogenetic stimulus conditions (light on/hyperpolarized, light off/baseline) were interleaved to avoid long-term adaptation effects.

#### Analytical Methods

Spike sorting for single channel recordings was performed online and validated offline using Waveclus (Quiroga et al., 2004). Spike sorting for multichannel electrodes was performed offline using the KlustaKwik software suite (Rossant et al., 2015). Isolation of the unit was confirmed by the waveform amplitude (absolute and relative to the background noise >3) and the interspike-interval distributions (VPm: mean of 0.22%, S1: mean of 0.38% of spikes in absolute refractory period of 1ms).

Although spike-triggered analysis for exploration of feature selectivity has been utilized widely in the thalamocortical circuit of the visual (Butts et al., 2007; Jones and Palmer, 1987; Lesica and Stanley, 2004; Reid and Alonso, 1995) and auditory (Eggermont et al., 1983; Theunissen et al., 2000) pathway, the application of this approach in the somatosensory pathway has been limited to a fairly small number of studies across thalamus (Petersen et al., 2008) and cortex (Estebanez et al., 2012; Maravall et al., 2007; Ramirez et al., 2014). Here we implemented spike-triggered analysis across VPm thalamus and S1 cortical layer 4. Specifically, feature selectivity was first estimated for each recorded unit using a simple spike triggered average (STA) (Schwartz et al., 2006).

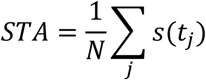

Where N is the number of spikes and *s* is the stimulus segment in a window surrounding each spike (−30 to +5 ms, spike-triggered ensemble). The burst and tonic triggered averages were computed from burst and tonic spikes, respectively. Burst spikes were classified here from the extracellular recordings as two or more spikes with an inter-spike interval of less than four milliseconds with the first spike in the burst preceded by 100 milliseconds of silence. The baseline/hyperpolarized condition triggered averages were computed from all spikes in a given stimulus condition. The bootstrap estimate of the confidence intervals on the spike triggered average was computed as the +/- 2 standard deviation of the shuffled STA distribution across 500 repetitions (Schwartz et al., 2006). Note that we implemented multiple techniques of estimating the feature selectivity of the neurons including spike triggered covariance, generalized linear models, and nonlinear-input models (McFarland et al., 2013). The results were qualitatively consistent across all methods employed, so we chose to use spike triggered average throughout the manuscript due to its simplicity.

The signal-to-noise ratio of the recovered STA was quantified as the peak-to-peak amplitude of the STA within 10 milliseconds of the spike (where the significant filter activity is contained) divided by the peak-to-peak amplitude of the STA from 30 to 20 milliseconds before the spike (where there is no expected filter information). An SNR value of 1 means the amplitude of the STA near the spike time is not different from the amplitude of the noise fluctuations. Therefore, any units with an SNR value less than 2 were excluded from further analysis.

There was significant diversity in the resultant STA structure across recorded neurons. Utilizing the approach of Estebanez et al. to make comparisons of the feature selectivity across the population of recorded neurons, we conducted a principle components analysis of the recovered STA (Estebanez et al., 2012). The first two principle components accounted for the majority of the variance (71.8% VPm, 78.4% S1), and were interpreted as representative of the primary structure in the sensory input relevant for spiking in the population.

The STA represents structure in the sensory input that is captured in the mean across spiking activity, and cannot capture any structure that may be in higher order statistical properties of the stimulus. Spike-triggered covariance (STC) can also be calculated as an alternative, as an attempt to uncover high-order structure in the sensory stimulus that is associated with neuronal firing (Estebanez et al., 2012). Note that while implemented in this analysis, STC analysis of both VPm and S1 neurons did not reveal significantly different structure for the neurons recorded under our experimental conditions, and are thus not shown here, and even more importantly, this analysis did not reveal a shift in feature selectivity across thalamic states.

The non-linearity (*P*(*spike*|*y*)) was estimated as the ratio of the probability of spike-triggered stimuli (*P*(*y*|*spike*)) to the probability of any stimulus segment in the stimulus (*P*(*y*)) multiplied by the mean firing rate of the neuron(*P*(*spike*)) (Schwartz et al., 2006):

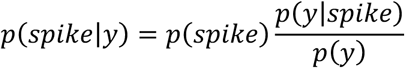

Where y is defined as the stimulus (s) convolved with the feature selectivity of the unit (STA) (Lesica et al., 2007), referred to as filtered stimulus. Because the slope of the static non-linearity is determined by the separation between the spike triggered ensemble and the Gaussian distributed white-noise, as the spike triggered ensemble distribution becomes more selective (i.e. the mean moves away from the filtered stimulus distribution), the separability of the distributions increases, and the slope of the non-linearity also increases. Intuitively, this means that the shape of the non-linearity gives an estimate of the separability of the spike triggered ensemble and the stimulus distribution, or how strongly tuned a neuron is for that particular feature, given by the STA. A steeper slope in the non-linearity suggests a stronger tuning than a shallower slope. Therefore, we also assessed the spiking nonlinearity as a function of the spike classification. For all conditions, the best estimate of the STA was defined as tonic spike triggered average for thalamic units and the baseline thalamic state (i.e. not optogenetically manipulated) for the cortical units. Throughout the manuscript, we separate the firing rate (p(spike)) from the shape of the non-linearity (p(y|spike)/p(y)) to avoid confounding differences in firing rate with differences in tuning.

The precision in the noise evoked firing was estimated for each spike classification (tonic, burst, baseline, hyperpolarized). The precision was defined as:

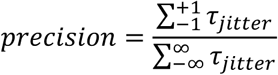

Or the number of spikes with τ_*jitter*_ values of +/- 1 ms duration normalized by the total number of spikes. τ_*jitter*_ is defined for each spike as the temporal lag (t_lag_) for the peak correlation between the STA and the stimulus segment surrounding that spike (−30 ms to +5 ms).

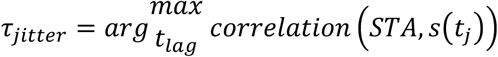

The τ_*jitter*_ distribution was normalized by the total number of spikes in each condition (tonic, burst, baseline, hyperpolarized).

All pairwise statistical comparisons were computed using a Wilcoxon signed rank test unless otherwise noted.

## Results

### Spike triggered analysis in the thalamus and cortex

We recorded thalamic and cortical extracellular spiking activity in response to sensory white-noise stimulation of a single whisker in the vibrissa pathway of the fentanyl-cocktail anesthetized rat to enable long-duration, controlled measurements needed for precise estimates of feature selectivity (Figure 1A, see Methods). We estimated the feature selectivity for each unit as the spike triggered average (STA), which captures the features of the sensory stimulus that tended to precede spiking, and the static, point nonlinearity, which captures the translation into suprathreshold spiking activity (Figure 1B; see methods). Although this quantification was performed on longer unique noise segments, we also recorded the response to short (4-10 second) frozen white-noise segments to examine the response across trials. Figure 1C shows an example recording from a simultaneously recorded pair of neurons in topographically aligned regions of the thalamus (left column, ventral posteromedial nucleus, VPm) and cortex (right column, primary somatosensory cortex, S1) in response to the repeated presentation of a single frozen white-noise segment (top of each column). Across trials, the repeatability of the response to the noise stimulus is apparent in the raster plot, with clear vertical patterns across trials. Spike-triggered analysis has been widely utilized in studying feature selectivity in the visual (Butts et al., 2007; Jones and Palmer, 1987; Lesica and Stanley, 2004; Reid and Alonso, 1995) and auditory (Eggermont et al., 1983; Theunissen et al., 2000) pathway, but has been utilized in the somatosensory pathway in only a relatively small number of studies (Estebanez et al., 2012; Maravall et al., 2007; Petersen et al., 2008; Ramirez et al., 2014). Here, to explore the feature selectivity of these recorded neurons, the spike-triggered average (STA) was computed for the thalamic and cortical unit for stimulus segments from −30 milliseconds prior to the spike to +5 milliseconds afterwards at a 0.2ms resolution (Figure 1C, bottom). The VPm STA shows clear feature selectivity in the 10-15 milliseconds prior to the thalamic spike as evidenced by the large amplitude of the STA relative to the shuffled case (grey confidence intervals). Beyond 15 milliseconds prior to the spike, the VPm STA is essentially flat and within the confidence bounds on the shuffled process. This suggests that, on average, the thalamic unit is only sensitive to the stimulus occurring in the previous 10-15 milliseconds. The S1 unit also displays feature selectivity as evidenced by the shape and amplitude of the S1 STA immediately prior to the cortical spike relative to the shuffled case. Although the VPm STA is nearly ten times as large in amplitude as the S1 STA, the similarity in the temporal dynamics can be visualized by shifting the VPm STA by 2 milliseconds relative to the S1 STA (Figure 1C, bottom, S1 STA black, VPm STA shifted by 2 milliseconds and scaled by a factor of 0.1 as grey dashed line).

**Figure 1:**
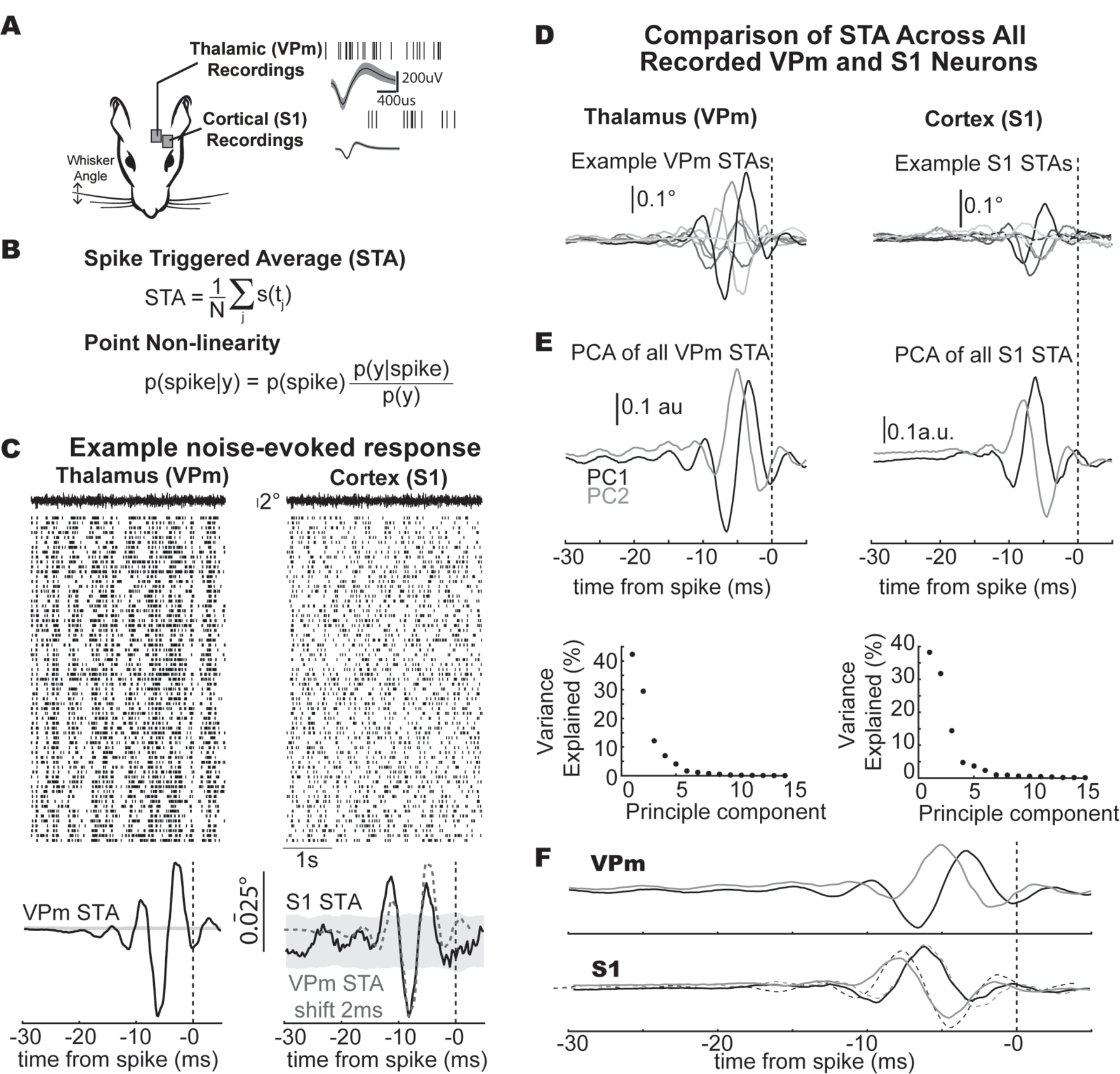
Spike-triggered analysis in the thalamocortical circuit of the rat whisker pathway. A. Experimental paradigm. B. Feature selectivity is assessed from the neural response to sensory white-noise as the spike triggered average and the associated point nonlinearity. C. Top: Example noise evoked response from simultaneously recorded topographically aligned pair of neurons. Bottom: Recovered STA for the example units. The black dotted vertical line indicates the time of the spike. D. Representative example thalamic and cortical STA (grey shades indicate different units). E. First two principle components computed using principle component analysis (PCA) of all recovered STA for thalamic neurons (n = 30) and cortical neurons (n = 32). F. Overlay of cortical STA PCs and temporally shifted thalamic PCs (shift of 2.5ms).

While this simple comparison provides an interesting observation for a single pair of topographically aligned neurons, we also made comparisons of the feature selectivity across the population of recorded STAs for thalamus and cortex. First, we visualized the shape of the spike triggered average for a sample of example thalamic (Figure 1D left; greyscale) and cortical (Figure 1D right; greyscale) units. These STA filters cannot be simply averaged together to give an estimate of the population filter due to differences in the phase and directionality of the recovered STA across different recorded units. Instead, utilizing an approach from Estebanez et al., we performed a principle component analysis on the set of recovered thalamic and cortical STA filters across recordings to identify salient filter properties that generalized (Figure 1E, Estebanez et al., 2012). The first two principle components for the spike triggered averages of both thalamus and cortex explain the majority of the variance for the set of recovered filters, similar to what has been seen previously for cortex (Estebanez et al., 2012). Furthermore, a simple time shift of 2.5 milliseconds for the VPm principle components relative to the S1 principle components (Figure 1F, dashed lines) demonstrates the similarity in the STA subspace spanned by these principle components. It seems that despite not necessarily being recorded simultaneously or even in the same animal, there is a high degree of overlap in the low dimensional subspace of feature selectivity for thalamocortical neurons in the whisker pathway. Note that we further analyzed the spike triggered covariance (STC) for the same data and found that the subspace spanned by the recovered linear filters matched the subspace estimated using the linear filters recovered using STA, thus not revealing any higher-order feature selectivity in the data.

### Tonic and burst spike triggered analysis in thalamus

Inherent in the spike triggered analysis, however, is an assumption that the average filter is representative of the sensory stimulus preceding all spikes (Stanley, 2002). Yet neurons in the thalamus are well known for exhibiting two fundamentally different types of firing: tonic spiking and burst firing mediated through T-type calcium channels (Suzuki and Rogawski, 1989). Burst spikes were classified here from the extracellular recordings as two or more spikes with an inter-spike interval of less than four milliseconds with the first spike in the burst preceded by 100 milliseconds of silence (Figure 2A, see methods). Using this classification, we asked if or how the feature selectivity of an individual thalamic unit changes as a function of the spiking mechanism in the whisker pathway.

**Figure 2:**
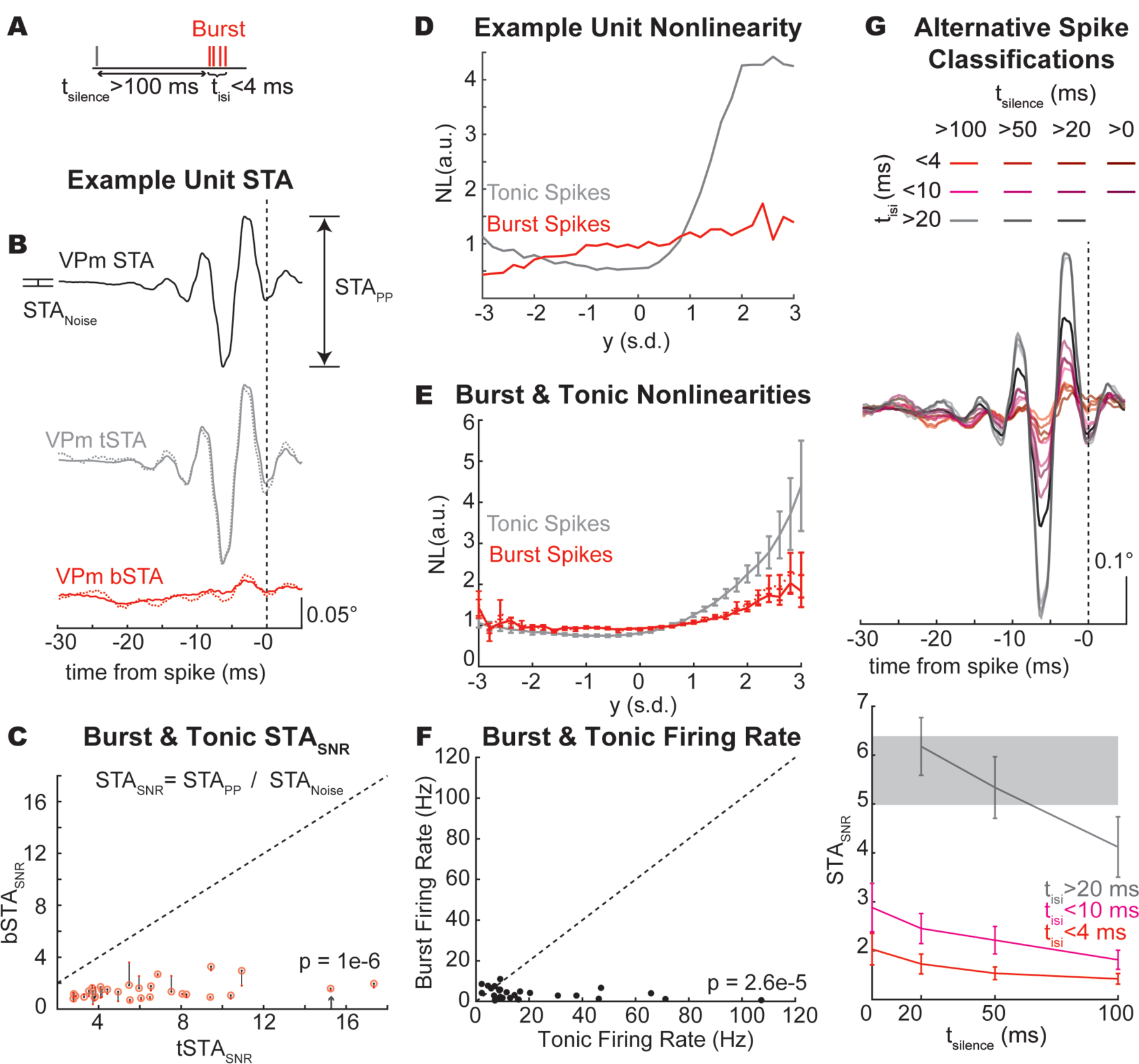
Spike triggered analysis of burst and tonic spikes in thalamic neurons. A. Burst definition. B. STA from thalamic unit presented in Figure 1 estimated from all spikes (black solid line, n = 44105 spikes), tonic spikes (tSTA; grey solid line, n = 36558 spikes), a subsample of tonic spikes (tSTA; dashed line, n = 2363 of 36558 spikes) all burst spikes (bSTA; red solid line, n = 7547 spikes), and the first spike in the burst (bSTA; red dashed line, n = 2363 spikes). C. STA_SNR_ across recorded population (n = 30 units). Black dots depict the STA_SNR_ for all bSTA computed from all burst spikes while red circles indicate the STA_SNR_ for bSTA computed from only the first spike in the burst (p = 1e-6). Arrow depicts example unit in A. D. Example unit tonic and burst spike nonlinearity. E. Average non-linearity across all units. F. Burst and tonic firing rate (p(spike)) across recorded population (p = 2.6e-5). G. Spike triggered average using different spike classifications (Bottom plot: n = 30 thalamic units).

In the thalamic recordings, tonic and burst spikes were interspersed throughout most of the recordings. For the example thalamic unit presented in Figure 1B, we computed the spike triggered average from all spikes (STA), the tonic spike triggered average from only tonic spikes (tSTA), and the burst spike triggered average from only spikes that are classified as being part of a burst (bSTA) (Figure 2B). The tSTA (grey) closely resembles the STA computed from all spikes (black) while the bSTA (red) is significantly degraded as evidenced by the flat shape of the filter. To compare the difference between burst and tonic feature selectivity across thalamic units, we quantified the signal-to-noise ratio of the STA (STA_SNR_, see methods). Across all thalamic units, the STA_SNR_ was higher for tonic spikes (tSTA_SNR_) than for burst spikes (bSTA_SNR_) (Figure 2C). Note that again we calculated the STC under these conditions and found the same result that the STC_SNR_ was higher for tonic spikes than for burst spikes (p = 2e-4), and that there was not a fundamental shift in representation between tonic and burst spiking.

If the timing of spikes within a burst is not repeatable and structured, the presence of these additional spikes will serve to destroy the temporal structure in the feature selectivity as revealed by the spike triggered analysis. When the bSTA was computed from only the first spike in each burst (Figure 2B, red-dashed line), there was no apparent feature selectivity for this example unit. This can also be visualized across units in the STA_SNR_ where the bSTA_SNR_ is plotted when computed from all burst spikes (black dot) and when computed from the first spike in each burst (red circle, Figure 2C). Therefore, including all spikes in a burst (or not) does not strongly impact the ability to estimate the feature selectivity from the STA.

Given the estimated feature selectivity, we can compute the static non-linearity, or the input-output function, which provides a mapping between this filtered stimulus (y) and the spiking response of the neuron (p(spike|y)) by taking the ratio of the p(y|spike) to the p(y) (Figure 1B, see methods). Here, we used the tSTA as the filter for all spiking conditions when estimating the non-linearity. The probability of the filtered stimulus (p(y) remains unchanged when the filter is held constant. Therefore, any change in the non-linearity is then only due to changes in the probability of the filtered stimulus given that a spike occurred (p(y|spike)), or the spike triggered ensemble. In this example unit, we found that the tonic spikes were well tuned to the STA, as evidenced by the steep slope of the non-linearity while the burst spikes were not well tuned to the STA, as evidenced by the relatively flat non-linearity (Figure 2D). This trend was consistent across units where the burst spikes showed reduced tuning to the STA as compared to tonic spikes as assessed by the slope of the spiking nonlinearity (Figure 2E). Here, we have separated the difference in the slope of the non-linearity from the difference in the prevalence of burst and tonic spikes (p(spike) or firing rate), which is markedly higher for tonic spikes than for burst spikes (Figure 2F). Furthermore, we tested alternative burst spike classifications and quantified the implication for the STA (Figure 2G, top). Across spiking classifications, increased periods of silence prior to the spike (t_silence_) led to decreased STA_SNR_ while bursts of spikes (t_isi_<4 or <10) had consistently lower STA_SNR_ relative to tonic spikes (t_isi_>20) (Figure 2G, example unit in middle, population data in bottom). This suggests that any tonic spiking incorrectly classified as part of a burst would serve to increase the amplitude, and thus the SNR, of the burst-triggered average, but the conservative definition of a burst pattern that we have used here likely minimizes this effect. Therefore, our data suggests a reduction in stimulus selectivity for burst spiking perhaps not due to a true loss of feature selectivity but instead due to the analytical framework which requires neurons having a high temporal precision relative to the timescale of the stimulus features.

### Optogenetic manipulation of thalamic state

The previous analysis was conducted by presenting sensory white-noise stimuli and parsing measured thalamic spiking activity into tonic and burst classes, while these classes of spiking were intermingled throughout the recordings. However, the thalamus was in tonic firing mode, with relatively low burst firing rates (Figure 2F). Here, we used optogenetic hyperpolarization of the thalamic neurons not to silence the thalamic neurons, but instead to shift the thalamus into a burst firing mode during sensory white-noise stimulation (Figure 3A). Using this optogenetic manipulation, we asked whether the optogenetically manipulated firing mode (baseline and hyperpolarized conditions) of the thalamus impacts encoding and how this relates to the classified burst/tonic modes.

**Figure 3:**
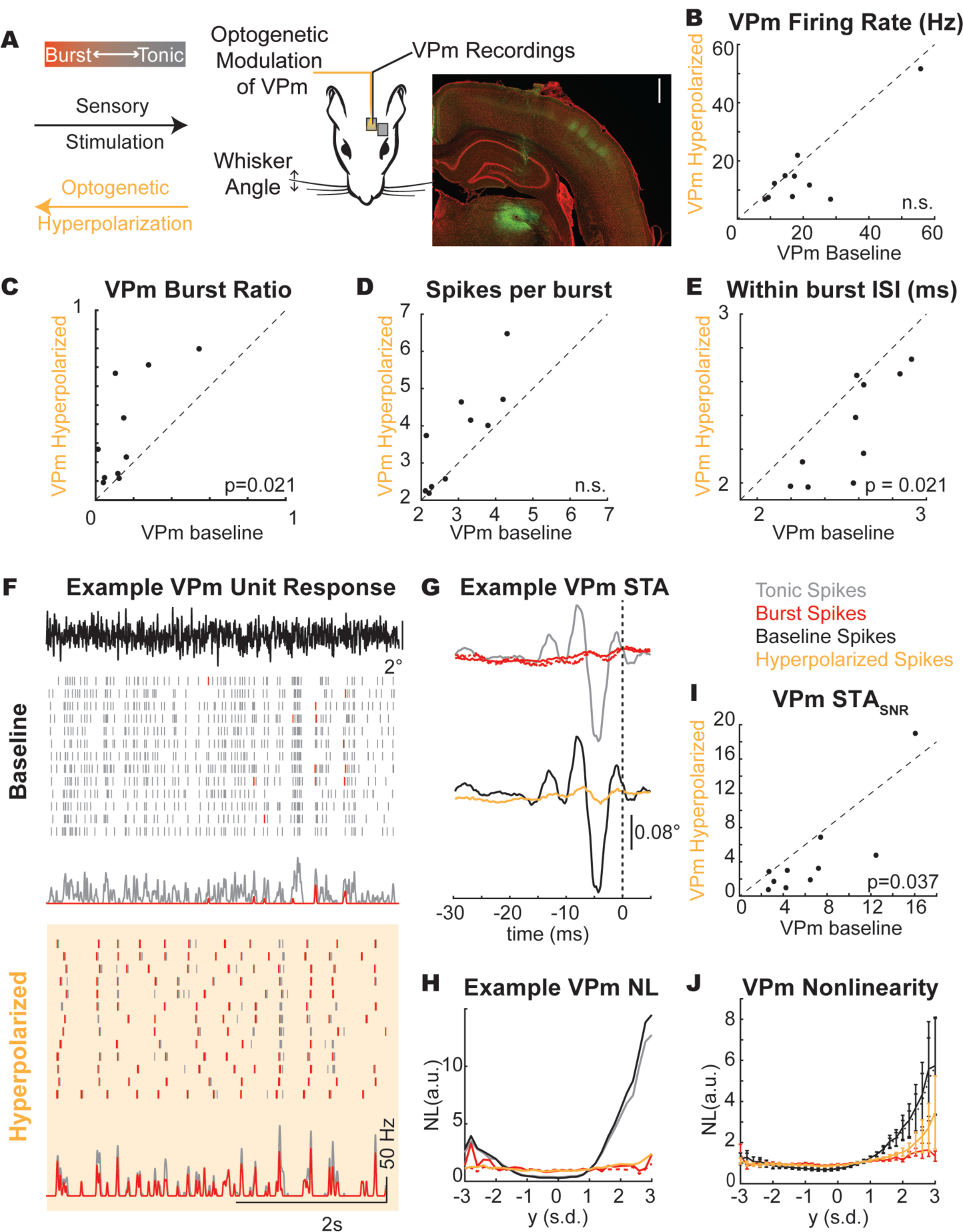
Optogenetic Manipulation of Thalamic State. A. Experimental paradigm. Scale bar: 1 mm. YFP signal (green) shows injection site in thalamus and axonal projections in cortex. B. Baseline and hyperpolarized firing rate (p(spike)) across recorded population (n = 10 units. p>0.05). C. Characterization of the burst ratio (p = 0.021), D. the spikes per burst (p>0.05), and E. the within burst ISI (p=0.021) during baseline and hyperpolarized conditions across the population of thalamic units (n = 10 units). F. Example thalamic response to frozen white-noise segments without (baseline) and with (hyperpolarized) optogenetic stimulation. Tick marks in raster plot are pseudocolored to demonstrate classification as burst (red tick) or tonic (grey tick) spikes. G. Spike triggered average computed as a function of spike classification (burst/red, first spike in burst/red dashed, tonic/grey) or thalamic state (baseline/black, hyperpolarized/yellow). H. Example unit tonic/burst and baseline/hyperpolarized spike nonlinearity. I. STA_SNR_ across recorded population (n = 10 units). J. Average non-linearity across all units (n = 10 units).

Here, we recorded the thalamic response to sensory white-noise with and without the presence of a light stimulus (hyperpolarized and baseline conditions, respectively). We found no significant change in the thalamic firing rate between hyperpolarized and baseline conditions (Figure 3B), but the firing mode of the thalamus did shift towards burst firing (baseline burst ratio = 0.16 ± 0.15, hyperpolarized burst ratio = 0.36 ± 0.27, n = 10 units, Figure 3C). Interestingly, the mean firing rate in VPm in response to sensory white-noise was not significantly different between the baseline and hyperpolarized conditions, reflecting a “replacement” of tonic firing with burst firing. It is also is apparent that the bursts in the baseline and hyperpolarized conditions are not occurring at similar times, despite a common sensory drive, suggesting some differences in the stimulus properties to which the bursts are sensitive across the two conditions. Bursts in the hyperpolarized condition showed a similar number of spikes per burst (Figure 3D) with a shorter inter-spike interval within a burst (Figure 3E). For an example unit, we have plotted the spiking response to a frozen white-noise segment without optogenetic stimulation (Figure 3F, baseline condition) and with optogenetic stimulation (Figure 3F, hyperpolarized condition). We have pseudo-colored the tonic spikes grey and the burst spikes red to qualitatively visualize the thalamic firing mode (Figure 3F). In the baseline condition, the response is primarily tonic as evidenced by the grey raster plots (Figure 3F, Baseline, BR = 0.10). In the hyperpolarized condition (optically stimulated), the firing mode is biased towards a burst encoding scheme, as evidenced by the prevalence of red burst spikes (Figure 3F, Hyperpolarized, BR = 0.67). The tonic STA showed pronounced feature selectivity for this unit while the burst STA did not (Figure 3G top), consistent with the earlier findings reflective of the loss in timing precision for the first spike of a burst (Figure 2). In the optogenetically manipulated states, the baseline STA has prominent feature selectivity while the hyperpolarized condition is much smaller in amplitude (Figure 3G bottom). Qualitatively, we can see that the STA from the hyperpolarized condition reflected the STA obtained from the burst spiking in the previous analysis. Again, a subsequent analysis of the STC revealed representations that were redundant with the spike-triggered average, and no apparent shift in feature selectivity between the tonic and burst firing modes.

The similarity between the burst spike response and hyperpolarized condition can also be seen in this example nonlinearity where the burst and hyperpolarized nonlinearities are effectively flat while the tonic spikes and baseline condition show obvious tuning (Figure 3H). Across units, we found an overall reduction in the STA_SNR_ for the hyperpolarized condition relative to the baseline condition (Figure 3I, p = 0.037). We also found that the tuning of the nonlinearity was lower for the hyperpolarized condition relative to the baseline condition as reflected in the overall gain/slope (Figure 3J). Importantly, the baseline and hyperpolarized conditions both contain burst and tonic spikes. Instead of completely separating the firing modes into all burst spikes or all tonic spikes, we have optogenetically altered the spiking probabilities such that the baseline condition has more tonic spikes and the hyperpolarized condition has more burst spikes (Figure 3C) while maintaining similar numbers of spikes (Figure 3B). The similarities between the STA and the NL properties of the burst and hyperpolarized state as well as the tonic and baseline state suggest that there was no discernable difference for the estimation of feature selectivity when assessed based on the state of the thalamus at the time of the stimulus (hyperpolarized/baseline) versus the spike type classification (burst/tonic).

### Temporal precision of thalamic firing modes

Given the difference between the recovered estimates of burst/hyperpolarized and tonic/baseline feature selectivity and the implications for timing precision in the thalamocortical circuit, we implemented a series of computational controls to identify any potential shortcomings of the methodologies that could underlie these results.

The first issue we considered was the overall difference in spike rates. Spike triggered analyses require a large number of spikes to effectively estimate the underlying selectivity. The proportion of spikes classified as bursts was lower than the spikes classified as tonic (Figure 2F) as quantified by the burst and tonic firing rate. Therefore, it was possible that we could not recover an STA for the burst spike condition due to the reduced number of burst spikes relative to tonic spikes. In an example unit, we computed the tSTA using only a subset of the spikes (n = 2363 of 36558 spikes corresponding to n = 2363 bursts with n = 7547 burst spikes) and found that the linear filter was essentially identical to the tSTA (Figure 2B, grey dashed line). We computed this for all thalamic units and again found that the burst-count matched tSTA was also significantly larger than the bSTA. Furthermore, there was no statistically significant difference in the firing rate between the baseline and hyperpolarized optogenetic conditions (Figure 3G), but still the difference in the STA persisted. This suggests that simple spike counts alone were insufficient to explain the difference in the tonic/baseline STA and the burst/hyperpolarized STA.

The second issue we considered was the inherent assumption that the feature selectivity for each unit could be recovered as the STA. It was possible that the burst STA was not recoverable because the burst firing mode was better estimated by a symmetric nonlinearity and therefore the filter could only be recovered using spike triggered covariance (STC) techniques. The STC approach has been previously implemented in the vibrissa pathway (Estebanez et al., 2012; Maravall et al., 2007; Petersen et al., 2008), revealing potential feature selectivity not captured with STA in some conditions, and was therefore important to consider here. We therefore computed the STC for all recorded thalamic units and compared this for each spiking condition. Although the dataset was more limited because the number of units with a significant STC filter was lower than those with a significant STA filter (n = 13 units with STC filter compared to n = 30 units with STA filter), the same trends regarding the reduction in the amplitude of the filter (STC_SNR_) and the slope of the symmetric nonlinearity persisted (as described in results). Therefore, this suggests that the method of extracting the feature selectivity (STA compared to STC) was insufficient to explain the inability to estimate the feature selectivity in the hyperpolarized/burst spiking conditions. However, it is also possible that the feature selectivity for a given neuron shifts to a higher-order space as firing modes transition from tonic to burst firing. If the burst firing could be associated with higher-order structure in the sensory stimulus, it may only be revealed using STC analysis. We thus conducted a STC analysis of the recorded neurons. First, we found that the tonic STA_SNR_ was significantly larger than the burst STC_SNR_ (p = 3e-6), suggesting that the feature selectivity did not simply shift from the first order estimate of the STA to higher order representations captured by the STC. Then we assessed the probability that a unit with a significant filter, as assessed using STA, also showed a significant filter, as assessed using STC. We found that all 13 units with significant filters also showed significant STA filters. This emphasizes the complexity of the stimulus representation, but further underscores that the stimulus representation did not simply shift from tonic to burst firing. Therefore, this analysis revealed that in general the high-order structure captured by STC was insignificant compared to the first order structure revealed by STA, and that when there was a loss of structure in the STA in the burst mode of firing, no new higher-order feature selectivity emerged through the STC analysis.

The third assumption made throughout the analysis was that burst spikes are actually driven by sensory stimuli such that there is a recoverable burst spike feature selectivity. The alternative explanation would be that burst spikes are not feature selective and instead occur randomly due to intrinsic or other non-sensory processes. To assess this, we quantified the trial-to-trial repeatability for bursts in response to frozen white-noise segments. As can be seen in Figure 3B, the qualitative assessment of temporally aligned bursts in response to the frozen white-noise segment suggests that the bursts are driven by the sensory stimulus in a repeatable way. For units with a sufficient number of repeated trials, we computed the reliability of the burst spiking as the correlation between the peristimulus time histogram of even and odd trials in response to the frozen white-noise segment. We found that all units showed greater reliability than what is expected based on just the temporal correlations in the burst spiking (shuffle control, p = 0.002, n = 10 thalamic units). This suggests that the bursts are not randomly generated or due entirely to a non-stimulus related phenomenon.

From these controls, we propose that the difference in the spike triggered encoding properties could not be attributed to differences in the overall spike rates, the temporal properties of the spikes within the burst, or the mechanism of filter estimation. Instead, we propose that the burst spikes are driven by the sensory stimulus and have an underlying feature selectivity, but that this cannot be recovered using spike triggered techniques due to the reduced temporal precision of burst spiking relative to tonic spikes.

Recovering feature selectivity from spike-triggered analysis relies on precise temporal spiking relative to the sensory stimulus. To simulate degradation of the spike timing precision, we added independent samples of normally distributed temporal jitter of varying amplitudes (standard deviation of the jitter distribution) to each tonic spike for an example unit and computed the STA (Figure 4A). Across units, we quantified the degradation of the STA as the jittered-STA_SNR_ normalized by the tSTA_SNR_ (0 ms jitter). The jittered-STA_SNR_ (black) is within the band expected for the bSTA_SNR_ with the addition of 4 milliseconds of jitter to the spike times (red shaded, Figure 4A, right). We propose that the effects of temporal jitter are particularly evident for whisker selectivity, presumably due to the short temporal duration of the filters (approximately 10-15 milliseconds in duration, Figure 1F).

**Figure 4:**
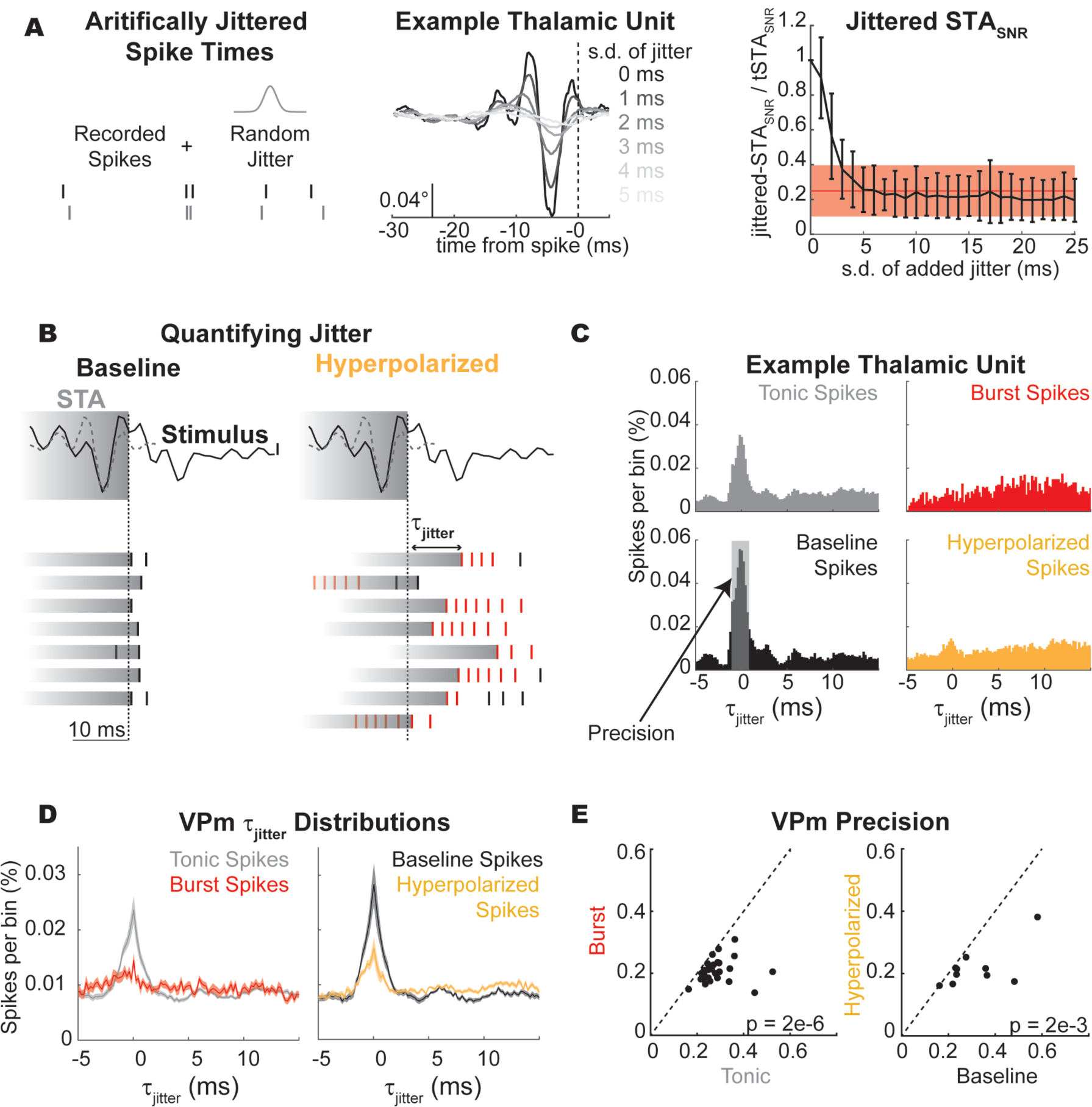
Thalamic timing variability in the response to the sensory white-noise. A. Effect of increased jitter on thalamic STA (schematic: left, example: center). Across units (n = 30), the normalized amplitude of the jittered STA (jitteredSTA_SNR_/tSTA_SNR_) was plotted across jitter intensities (black, right). The normalized amplitude of the burst STA (bSTA_SNR_/tSTA_SNR_) is shown in red (mean ± s.d.). B. In this example, the same stimulus (black; scale bar: 0.1°) was presented with and without optogenetic hyperpolarization (data from example unit presented in Figure 3 with associated STA (grey dotted line; scale bar: 0.025°)). C. Example unit τ_jitter_ distributions for burst (n = 2107 spikes), tonic (n = 23023 spikes), baseline (n = 11361 spikes), and hyperpolarized conditions (n = 11662 spikes). D. Average τ_jitter_ distributions for burst/tonic spikes (n = 30 units) and baseline/hyperpolarized condition (n = 10 units). (mean +/- sem). E. Precision for burst/tonic spikes (n = 30 units, p = 1e-6) and baseline/hyperpolarized condition (n = 10 units). (mean +/- sem, p=1e-3).

Given the marked effects of jitter on the ability to recover the STA, we investigated the variability in the spike timing relative to the noise stimulus (Figure 4B). For this example unit, we have identified a segment in the noise stimulus that closely resembles the tonic STA for this unit and elicits a reliable spiking response (Figure 4B, top; stimulus – black, tSTA – grey dashed). The vertical dashed line indicates the spike time for the spike triggered average (t_0_) with the grey bar indicating the duration of the STA. If there was no variability in the neural spiking, the raster plots would all be perfectly aligned to t0 because the similarity between the stimulus and the STA would predict a spiking response at that time point. However, the timing of evoked neural responses is always variable to some extent and this can be visualized for this example response segment as the temporal variability of the spike times surrounding this stimulus feature in the noise stimulus (Figure 4B, as indicated by the grey stimulus bars that extend from the first spike response to this particular sensory feature). For this example snapshot, it is also apparent that the burst spikes in the hyperpolarized condition show greater temporal variability than the tonic spikes in the baseline condition.

To quantify this jitter across all spikes, we developed a τ_jitter_ metric that determines the time lag of the peak correlation between the STA and the stimulus segment (s(t_j_)) surrounding each spike (Figure 4C). Intuitively, this is a correlative method to identify the time lag between when we predict a spike is most likely to occur based on the STA and the stimulus (peak correlation) and when the spike actually occurred. For this analysis, we treated the tSTA as the true feature selectivity of the neuron across all spiking conditions because we could not recover a reliable estimate of the bSTA.

We computed τ_jitter_ for each spike and plotted τ_jitter_ distributions for each spike condition (tonic, burst, baseline, hyperpolarized). If a neuron was infinitely precise such that when the stimulus matched the spike triggered average, the neuron fired a spike without delay, this distribution would be represented by a delta function at τ_jitter_ equals zero. As the variability of the timing increases, the width of this distribution will also increase. For the tonic and baseline condition spikes, we found a clear peak in τ_jitter_ values at τ_jitter_ equals zero (Figure 4C, grey, black). For the burst and hyperpolarized condition spikes, we observe little-to-no peak in the τ_jitter_ metric at zero (Figure 4C, red, yellow). We computed the τ_jitter_ distribution across all thalamic units and found that the tonic and baseline spikes had higher peaks at τ_jitter_ equals zero than the burst and hyperpolarized spike conditions (Figure 4D). We quantified this statistically by computing a precision metric (Figure 4C) that computes the proportion of spikes within ±1 millisecond of τ_jitter_ equals zero (Figure 4E). The tonic and baseline spike condition were more precise than burst and hyperpolarized conditions.

These data suggest that tonic spikes showed greater temporal precision in response to the sensory white-noise than burst spikes and that this could underlie the difference in the recoverability of the feature selectivity in the thalamus between firing modes. It is well established that the timing of sensory inputs is particularly important in the thalamocortical circuit such that changes in thalamic spike timing could have large impacts on the downstream representation of sensory information in the cortex. Next, we investigated how these changes in temporal precision in optogenetically modulated thalamic states impact cortical encoding properties.

### Optogenetic modulation of thalamic firing modes directly impacts cortical representation of sensory information

Cortical neurons that receive direct thalamic input are integrating information over a population of thalamocortical neurons that can be exhibiting different firing characteristics. This makes it difficult to determine the impact of a single burst from a single neuron on information representation in the pathway. Instead, we used the optogenetic manipulation of thalamic state as presented in Figure 3 to bias the activity of the thalamic population towards burst firing (hyperpolarized condition) while recording the cortical activity extracellularly (Figure 5A).

**Figure 5:**
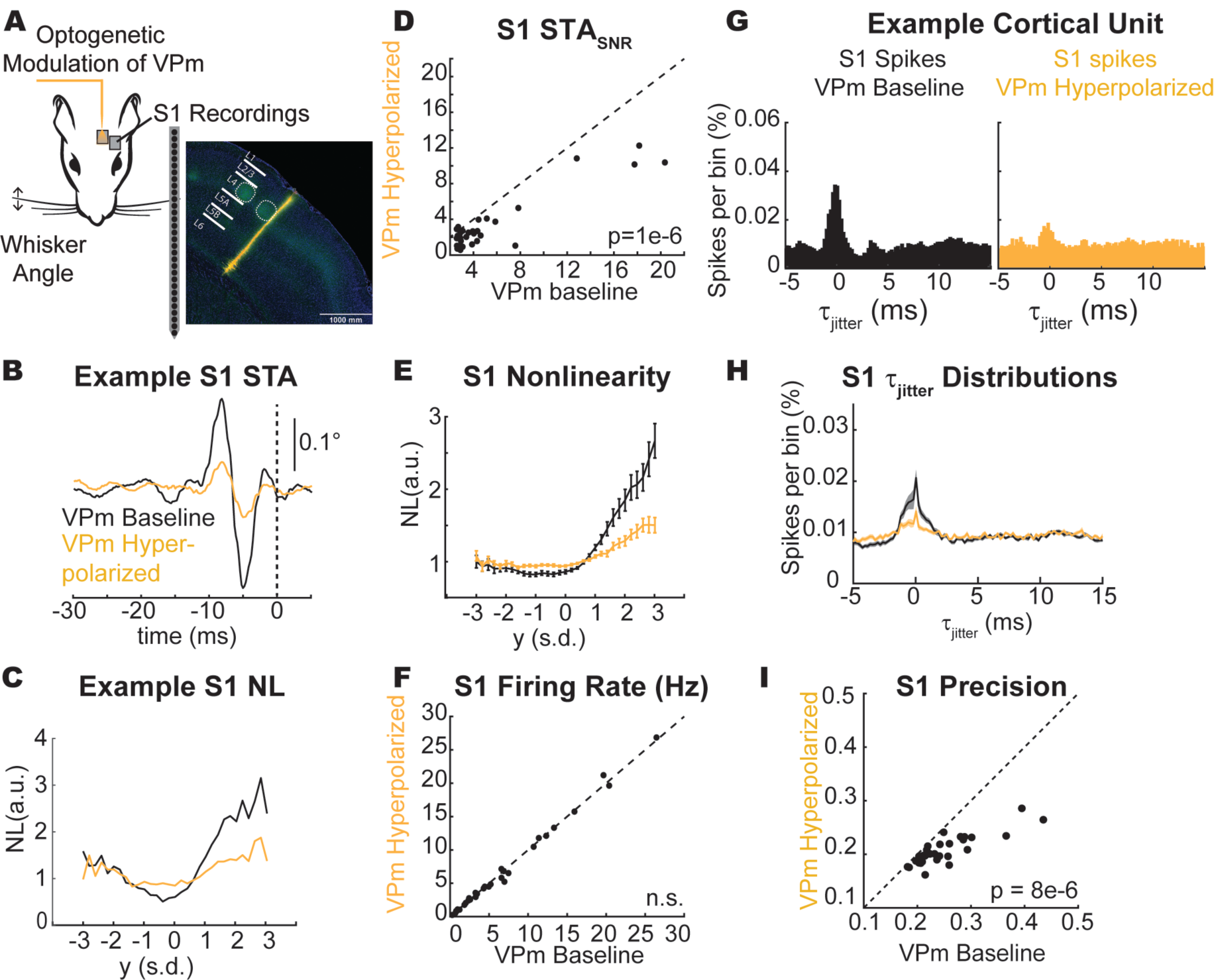
Optogenetic modulation of thalamic firing modes directly impacts cortical representation of sensory information. A. Experimental paradigm. Scale bar: 1 mm. YFP signal (green) shows axonal projections into cortex. DiI stain was used on the probe to confirm recording location. B. Example STA for a cortical unit recorded during optogenetic manipulation of the thalamus (black: baseline thalamic condition, yellow: hyperpolarized thalamic condition). C. Example cortical nonlinearity as a function of thalamic state. D. STA_SNR_ across cortical units (n = 32 units. p = 1e-6). E. Spiking nonlinearity across cortical units during thalamic manipulation (n = 32 units). F. Cortical firing rate during thalamic manipulation (n = 32 units, p = 0.31). G. Example cortical unit τ_jitter_ distributions for baseline (n = 12862 spikes) and hyperpolarized (n = 11848 spikes) thalamic conditions. H. Average τ_jitter_ distributions for baseline/hyperpolarized thalamic condition (n = 32 units, mean +/- sem). I. Precision for cortical spikes during baseline/hyperpolarized thalamic condition (n = 32 units, mean +/- sem, p=8e-6).

For an example unit, we have plotted the cortical STA in the baseline and hyperpolarized VPm conditions (Figure 5B). Here, the amplitude of the cortical STA was smaller when the thalamus is hyperpolarized compared to when it is not (Figure 5B). This cortical unit also shows a reduced tuning to the STA when the thalamus was hyperpolarized (Figure 5C). Across the population of recorded cortical neurons, the same effect seen in this example neuron of a reduced STA_SNR_ when the VPm was hyperpolarized compared to when it was not (Figure 5D) and a reduction in the tuning was present across all cortical units as quantified by the spiking nonlinearity (Figure 5E). These findings mirror what was seen for thalamic neurons when comparing the baseline and the optogenetically manipulated conditions demonstrating that the changes in thalamic encoding properties are propagated to cortex. Note that we again conducted the STC analysis with the S1 neurons and found the same reduction in STC_SNR_ when the VPm was hyperpolarized compared to when it was not (p = 9e-5), suggesting that the loss of feature selectivity in S1 with thalamic hyperpolarization was not just due to the transfer of feature selectivity to higher-order characteristics.

Interestingly, there was no significant difference in the sensory white-noise-evoked firing rate in the cortex as a function of the VPm condition (Figure 5F). This suggests that it was not overall spike counts influencing the cortical feature selectivity. Instead, we propose the temporal jitter in the thalamic spiking patterns propagated to cortex. We investigated the temporal precision of the cortical spiking in response to the sensory white-noise using the same methodology employed for the thalamus. As we saw for the thalamus, the cortical spikes from this example unit also showed greater temporal precision in response to white-noise whisker stimulation in the baseline VPm condition compared to the hyperpolarized VPm condition (Figure 5G) as evidenced by the peak in the τ_jitter_ distribution around τ_jitter_ equals zero. This effect was consistent across the population of recorded cortical units (Figure 5H) and showed significant differences in the precision of the cortical firing (Figure 5I). This suggests that the temporal jitter present in the thalamus is transmitted to the cortex where it also impacts the representation of sensory information.

## Discussion

Although there have been extensive investigations into the cortical state-dependent processing of the thalamocortical circuit, we know surprisingly little about how information is processed in a thalamic state-dependent manner. Here, using a combination of optogenetic manipulation and electrophysiological recording techniques, we have performed a series of experiments modulating the state of the thalamus (through constant optogenetic hyperpolarization) and quantified the effects on encoding in the thalamocortical circuit. Using this technique, we have coarse control of the firing mode in thalamus without altering the processing occurring from the whisker to thalamus, enabling us to decouple the changes in thalamic firing mode on thalamocortical processing from changes occurring in subthalamic processing. We found that, unlike the visual pathway, the feature selectivity of burst spikes in the vibrissa pathway could not be recovered using spike triggered techniques due to increased burst spike timing variability (loss of timing precision) relative to tonic spike timing. Recordings from barrel cortex during optogenetic manipulation of thalamic state demonstrated a loss in the temporal precision of the cortical spiking that also led to a degradation of the recovered feature selectivity. This suggests that bursts in the whisker pathway are less precise than tonic spikes during ongoing weak sensory stimulation and that this loss of temporal precision is propagated to cortex, which could have implications for the integration of complex patterns of sensory inputs.

Although spike-triggered analysis has been widely applied in various pathways, there have been comparatively few studies of this nature in the vibrissa pathway despite the extensive utilization of this model system. In this study, we focused primarily on timing in the thalamocortical circuit, and used the spike-triggered analysis as a vehicle to probe this issue rather than uncovering novel aspects of feature selectivity related to whisker kinematics. Nevertheless, it is important to note the similarities and differences between these studies. Spike-triggered averaging of VPm neurons in a study from Petersen et al. revealed very similar feature selectivity to what we report for VPm in baseline conditions here, with spiking tending to be preceded by a very fast transient, biphasic whisker deflection (Petersen et al., 2008). Studies in cortex, however, reveal more complex properties. Although the basic feature selectivity that we uncovered for S1 neurons in baseline conditions using STA was very similar to what has been observed in a subset of recorded neurons in other cortical studies (Estebanez et al., 2012; Maravall et al., 2007), further analysis using STC as well as more complex whisker stimulation paradigms in these studies identified more complex encoding properties. It should be noted that we restricted analysis to cortical S1 neurons that exhibited significant feature selectivity in the baseline condition with STA, but also observed other cortical neurons that showed significant feature selectivity only using STC, consistent with these previous studies. For these neurons, the filters recovered using STC analysis for tonic spikes were lost when they were computed for burst spikes, as the STC analysis is also reliant on precise spike-timing relative to the sensory stimulus.

Specific to our findings here related to thalamic firing modes, it is theoretically possible that in the optogenetically induced thalamic burst mode, the feature selectivity is not lost, but instead transformed to a type of selectivity that is not captured through the simple characteristics of the spike-triggered averaging or a stimulus selectivity that is fundamentally different from the tonic spike feature selectivity. As described above, previous studies have utilized STC analysis to successfully uncover complex feature selectivity in the visual (Touryan et al., 2005) and somatosensory (Estebanez et al., 2012; Maravall et al., 2007) pathways. However, when we extended the analysis here to the spike-triggered covariance (STC), it was not the case that the timing changes were captured through covariance analysis. For both the thalamic VPm and cortical S1 recordings, there was no apparent shift in feature selectivity from first-order structure (STA) to higher-order structure (STC) with a change in thalamic firing mode or state. Units with significant filters in the tonic spiking condition did not show significant filters in the burst spiking condition, even when assessed using both STA and STC. As with the spike triggered analysis, further analysis to explore burst feature selectivity that is fundamentally different from the tonic feature selectivity could not be pursued due to the timing variability of the spiking. We cannot identify this selectivity without a mechanism to measure, and compensate for, the increased spike timing jitter, as this framework is inextricably linked to the timing precision with which neurons spike relative to the sensory input.

These results could be interpreted as consistent with the view that bursts are not representing detailed stimulus information. However, there is evidence that bursts may convey more information than the “all-or-none” presence or absence of a burst through inter-burst spike timing and the number of spikes per burst (Mease et al., 2017), suggesting a role of temporally precise burst firing in information representation. Furthermore, thalamic bursting can be temporally precise within and across neurons in response to high intensity whisker stimuli (Whitmire et al., 2016). Instead, we propose that the temporal precision of the thalamic firing is a function of both the state of the thalamus and the intensity of the sensory stimulus. It has previously been shown that the temporal precision of thalamic encoding increases with the intensity of the sensory stimulus (Desbordes et al., 2008; Whitmire et al., 2016) while here we have shown that the temporal precision of the thalamic firing decreases with sustained hyperpolarization, which would naturally have implications for what signals do and do not get conveyed through the relatively narrow cortical window of integration (Gabernet et al., 2005). These two competing factors would enable the burst firing mode to encode high amplitude stimuli in a temporally precise fashion while low amplitude stimuli, such as the sensory white-noise presented here, would not be able to overcome the variability present in the burst state.

There are multiple mechanisms that could underlie the reduced temporal precision in the burst firing mode including variability introduced by the slow dynamics of the calcium depolarization, increased variability in the time to reach threshold due to the prolonged hyperpolarization of the baseline polarization, as well as potential changes in the integration properties of the thalamic neurons. Furthermore, these mechanisms could occur independently such that the variability across neurons is uncorrelated or these mechanisms could be coordinated in some way to enable correlated variability across the thalamic population. Both coordinated and uncoordinated jitter would have a detrimental effect on the ability to recover the STA because either the spike timing would no longer be locked to the stimulus itself or the input to the cortex would be temporally imprecise. However, coordinated jitter would maintain the information about the stimulus while uncoordinated jitter would degrade the recoverability of the stimulus features with the spike-triggered approach. Future work is needed to investigate the jitter in the burst spiking across the population to determine whether or not the variability in the spike timing is coordinated across thalamic units in this context.

While we have primarily considered thalamic state-dependent encoding as a feedforward representation from thalamus to cortex, the highly interconnected thalamocortical circuitry shapes coding properties in both feedforward and feedback manner. Changes in thalamic activity impact cortical activity which then provides feedback to thalamus to further alter activity (Crandall et al., 2015; Mease et al., 2014; Poulet et al., 2012; Reinhold et al., 2015; Wimmer et al., 2015). It is possible for thalamus to influence cortical state and for the cortex to influence thalamic state, but how this plays out during natural behaviors is not yet known and must be decoupled using more sophisticated techniques such as closed-loop control of neural activity (Bolus et al., 2018; Newman et al., 2015).

Given the importance of thalamic spike timing precision within and across neurons in transmitting information downstream to cortex (Bruno and Sakmann, 2006; Wang et al., 2010), alterations to thalamic state can shape multiple properties of the spiking inputs to cortex. Here, we have shown that thalamic state directly impacts the spike timing precision in the thalamus and cortex. Manipulation of the thalamic state can also lead to changes in the stimulus evoked cortical dynamics (Whitmire et al., 2017) and spatiotemporal cortical activation (Borden et al., 2019). Alterations to the state of the thalamus, or the baseline membrane potential, provides a biophysical mechanism for the thalamus to gate information flow to cortex. Furthermore, this mechanism could be under both feedforward and feedback control. This sets the stage for a dynamic interaction between thalamic and cortical states to drive highly interactive patterns of neural activity, ultimately controlling the integration of afferent signaling that underlies sensory percepts.

## Acknowledgements

This work was supported by NIH National Institute of Neurological Disorders and Stroke Grants R01NS085447 and R01NS104928. CJW was supported by the NIH NRSA Pre-doctoral Fellowship (F31NS089412). Confocal imaging was performed at the Georgia Tech IBB Imaging Core. We thank Peter Y Borden, Aurélie Pala, Christian Waiblinger, and Caleb Wright for helpful comments on the data analysis and manuscript.

## References

Alitto HJ, Weyand T, Usrey WM. 2005. Distinct Properties of Stimulus-Evoked Bursts in the Lateral Geniculate Nucleus. J Neurosci 25:514–523. doi:10.1523/JNEUROSCI.3369-04.2005

Bolus MF, Willats AA, Whitmire CJ, Rozell CJ, Stanley GB. 2018. Design strategies for dynamic closed-loop optogenetic neurocontrol in vivo. J Neural Eng 15:026011. doi:10.1088/1741-2552/aaa506

Borden PY, Wright NC, Sederberg AJ, Waiblinger C, Haider B, Stanley GB. 2019. Thalamic modulation and the shaping of cortical sensory representations in the awake and anesthetized mouse2019 Neuroscience Meeting Planner. Chicago, IL: Society for Neuroscience. p. Program No. 221.19.

Bruno RM, Sakmann B. 2006. Cortex is driven by weak but synchronously active thalamocortical synapses. Science 312:1622–7. doi:10.1126/science.1124593

Butts D a, Weng C, Jin J, Yeh C-I, Lesica NA, Alonso J-M, Stanley GB. 2007. Temporal precision in the neural code and the timescales of natural vision. Nature 449:92–5. doi:10.1038/nature06105

Castro-alamancos MA. 2002. Properties of primary sensory (lemniscal) synapses in the ventrobasal thalamus and the relay of high-frequency sensory inputs. J Neurophysiol 946–953.

Crandall SR, Cruikshank SJ, Connors BW, Crandall SR, Cruikshank SJ, Connors BW. 2015. A Corticothalamic Switch: Controlling the Thalamus with Dynamic Synapses. Neuron 86:1–15. doi:10.1016/j.neuron.2015.03.040

Denning KS, Reinagel P. 2005. Visual Control of Burst Priming in the Anesthetized Lateral Geniculate Nucleus. J Neurosci 25:3531–3538. doi:10.1523/JNEUROSCI.4417-04.2005

Desbordes G, Jin J, Weng C, Lesica NA, Stanley GB, Alonso J. 2008. Timing precision in population coding of natural scenes in the early visual system. PLoS Biol 6:e324. doi:10.1371/journal.pbio.0060324

Eggermont JJ, Johannesma PIM, Aertsen AMH. 1983. Reverse correlation methods in auditory research. Q Rev Biophys 16:341–414. doi:10.1017/S0033583500005126

Estebanez L, Boustani S El, Destexhe A, Shulz DE. 2012. Correlated input reveals coexisting coding schemes in a sensory cortex. Nat Neurosci 15:1–14. doi:10.1038/nn.3258

Gabernet L, Jadhav SP, Feldman DE, Carandini M, Scanziani M. 2005. Somatosensory integration controlled by dynamic thalamocortical feed-forward inhibition. Neuron 48:315–27. doi:10.1016/j.neuron.2005.09.022

Guido W, Weyand T. 1995. Burst responses in thalamic relay cells of the awake behaving cat. J Neurophysiol 74:1782–1786.

Jones JP, Palmer LA. 1987. The two-dimensional spatial structure of simple receptive fields in cat striate cortex. J Neurophysiol 58:1187–211.

Lesica NA, Jin J, Weng C, Yeh C-I, Butts D a, Stanley GB, Alonso J-M. 2007. Adaptation to stimulus contrast and correlations during natural visual stimulation. Neuron 55:479–91. doi:10.1016/j.neuron.2007.07.013

Lesica NA, Stanley GB. 2004. Encoding of natural scene movies by tonic and burst spikes in the lateral geniculate nucleus. J Neurosci 24:10731–40. doi:10.1523/JNEUROSCI.3059-04.2004

Maravall M, Petersen RS, Fairhall AL, Arabzadeh E, Diamond ME. 2007. Shifts in coding properties and maintenance of information transmission during adaptation in barrel cortex. PLoS Biol 5:e19. doi:10.1371/journal.pbio.0050019

Martinez-Conde S, Macknik SL, Hubel DH. 2002. The function of bursts of spikes during visual fixation in the awake primate lateral geniculate nucleus and primary visual cortex. Proc Natl Acad Sci 99:13920–13925. doi:10.1073/pnas.212500599

McFarland JM, Cui Y, Butts D a. 2013. Inferring nonlinear neuronal computation based on physiologically plausible inputs. PLoS Comput Biol 9:e1003143. doi:10.1371/journal.pcbi.1003143

Mease RA, Krieger P, Groh A. 2014. Cortical control of adaptation and sensory relay mode in the thalamus. Proc Natl Acad Sci U S A 111:6798–803. doi:10.1073/pnas.1318665111

Mease RA, Kuner T, Fairhall AL, Groh A. 2017. Multiplexed Spike Coding and Adaptation in the Thalamus. Cell Rep 19:1130–1140. doi:10.1016/j.celrep.2017.04.050

Newman JP, Fong M-F, Millard DC, Whitmire CJ, Stanley GB, Potter SM. 2015. Optogenetic feedback control of neural activity. Elife 1–24. doi:10.7554/eLife.07192

Petersen RS, Brambilla M, Bale MR, Alenda A, Panzeri S, Montemurro M a, Maravall M. 2008. Diverse and temporally precise kinetic feature selectivity in the VPm thalamic nucleus. Neuron 60:890–903. doi:10.1016/j.neuron.2008.09.041

Poulet JFA, Fernandez LMJ, Crochet S, Petersen CCHH. 2012. Thalamic control of cortical states. Nat Neurosci 15:370–2. doi:10.1038/nn.3035

Quiroga RQ, Nadasdy Z, Ben-Shaul Y. 2004. Unsupervised spike detection and sorting with wavelets and superparamagnetic clustering. Neural Comput 16:1661–1687. doi:10.1162/089976604774201631

Ramirez A, Pnevmatikakis E, Merel J, Paninski L, Miller KD, Bruno RM. 2014. The spatiotemporal receptive fields of barrel cortex neurons revealed by reverse correlation of synaptic input. Nat Neurosci 17:866–875.

Reid RC, Alonso J-M. 1995. Specificity of monosynaptic connections from thalamus to visual cortex. Nature 378:281–4. doi:10.1038/378281a0

Reinagel P, Godwin D, Sherman SM, Koch C. 1999. Encoding of visual information by LGN bursts. J Neurophysiol 2558–2569.

Reinhold K, Lien AD, Scanziani M. 2015. Distinct recurrent versus afferent dynamics in cortical visual processing. Nat Neurosci 18. doi:10.1038/nn.4153

Rossant C, Kadir SN, Goodman DFM, Schulman J, Belluscio M, Buzsáki G, Harris KD, Hunter MLD, Saleem AB, Grosmark A, Denfield GH, Ecker ASS, Tolias AS, Solomon SG, Carandini M, Belluscio M, Denfield GH, Ecker ASS, Tolias AS, Solomon SG, Buzsáki G, Carandini M, Harris KD. 2015. Spike sorting for large, dense electrode arrays. Nat Neurosci 015198. doi:10.1038/nn.4268

Schwartz O, Pillow JW, Rust NC, Simoncelli EP. 2006. Spike-triggered neural characterization. J Vis 6:484–507. doi:10.1167/6.4.13

Sherman SM. 2001. A wake-up call from the thalamus. Nat Neurosci 4:344–6. doi:10.1038/85973

Stanley GB. 2002. Adaptive spatiotemporal receptive field estimation in the visual pathway. Neural Comput 14:2925–2946. doi:10.1162/089976602760805340

Suzuki S, Rogawski MA. 1989. T-type calcium channels mediate the transition between tonic and phasic firing in thalamic neurons. Proc Natl Acad Sci U S A 86:7228–7232. doi:10.1073/pnas.86.18.7228

Swadlow HA, Gusev AG. 2001. The impact of “bursting” thalamic impulses at a neocortical synapse. Nat Neurosci 4:402–8. doi:10.1038/86054

Theunissen FE, Sen K, Doupe AJJ. 2000. Spectral-temporal receptive fields of nonlinear auditory neurons obtained using natural sounds. J Neurosci 20:2315–2331.

Touryan J, Felsen G, Dan Y. 2005. Spatial structure of complex cell receptive fields measured with natural images. Neuron 45:781–91. doi:10.1016/j.neuron.2005.01.029

Wang Q, Webber RM, Stanley GB. 2010. Thalamic synchrony and the adaptive gating of information flow to cortex. Nat Neurosci 13:1534–41. doi:10.1038/nn.2670

Wang X, Wei Y, Vaingankar V, Wang Q, Koepsell K, Sommer FT, Hirsch J a. 2007. Feedforward Excitation and Inhibition Evoke Dual Modes of Firing in the Cat’s Visual Thalamus during Naturalistic Viewing. Neuron 55:465–478. doi:10.1016/j.neuron.2007.06.039

Whitmire CJ, Millard DC, Stanley GB. 2017. Thalamic state control of cortical paired-pulse dynamics. J Neurophysiol 117:jn 00415 2016. doi:10.1152/jn.00415.2016

Whitmire CJ, Waiblinger C, Schwarz C, Stanley GB. 2016. Information Coding through Adaptive Gating of Synchronized Thalamic Bursting. CellReports 14:1–13. doi:10.1016/j.celrep.2015.12.068

Wimmer RD, Schmitt LI, Davidson TJ, Nakajima M, Deisseroth K, Halassa MM. 2015. Thalamic control of sensory selection in divided attention. Nature. doi:10.1038/nature15398

Wolfart J, Debay D, Le Masson G, Destexhe A, Bal T. 2005. Synaptic background activity controls spike transfer from thalamus to cortex. Nat Neurosci 8:1760–7. doi:10.1038/nn1591

